# ^68^Ga-Bisphosphonates for the Imaging of Extraosseous Calcification by Positron Emission Tomography

**DOI:** 10.1101/2022.11.15.516425

**Authors:** George. P. Keeling, Friedrich Baark, Orestis L. Katsamenis, Jing Xue, Philip J. Blower, Sergio Bertazzo, Rafael T. M. de Rosales

## Abstract

Radiolabelled bisphosphonates (BPs) and [^18^F]NaF (^18^F-fluoride) are the two types of radiotracers available to image calcium mineral *in vivo* (e.g. bone), yet only [^18^F]NaF has been widely explored for the non-invasive molecular imaging of extraosseous calcification (EC) using the highly sensitive nuclear imaging technique positron emission tomography (PET). These two radiotracers bind calcium mineral deposits via different mechanisms, with BPs chelating to calcium ions and thus being non-selective, and [^18^F]NaF being selective for hydroxyapatite (HAp) which is the main component of bone mineral. Taking into account that the composition of EC has been reported to include a diverse range of non-HAp calcium minerals, we hypothesised that BPs may be more sensitive for imaging EC due to their ability to bind to both HAp and non-HAp deposits.

To test this hypothesis, we report a comparison between the ^68^Ga-labelled BP tracer [^68^Ga]Ga-THP-Pam and [^18^F]NaF for PET imaging in a rat model of EC that develops macro- and microcalcifications in several organs. The presence of macrocalcifications was identified using preclinical computed tomography (CT) and microcalcifications were identified using μCT-based 3D X-ray histology (XRH) on isolated organs *ex vivo*. The morphological and mineral analysis of individual calcified deposits was performed using scanning electron microscopy (SEM) and energy-dispersive X-ray spectroscopy (EDX). The PET imaging and *ex vivo* analysis results demonstrated that while both radiotracers behave similarly for bone imaging, the BP-based radiotracer [^68^Ga]Ga-THP-Pam was able to detect EC more sensitively in several organs in which the mineral composition departs from that of HAp. We conclude that BP-based PET radiotracers such as [^68^Ga]Ga-THP-Pam have a particular advantage for the sensitive imaging and early detection of EC by being able to detect a wider array of relevant calcium minerals *in vivo* than [^18^F]NaF, and should be evaluated clinically for this purpose.

## Introduction

Bisphosphonates (BPs) are a class of drug with high affinity for solid calcium minerals such as hydroxyapatite (HAp) [1], the primary inorganic component of bone tissue [2], and accumulate particularly in areas of high mineral turnover, such as bone metastases [1, 3]. This accumulation, in combination with their small size and characteristically fast blood clearance, has made BPs a mainstay of medical imaging of bone diseases since the 1970s, most commonly in the form of [^99m^Tc]Tc-MDP using gamma-scintigraphy or SPECT imaging [3-5].

Technological advances in positron emission tomography (PET) have ushered in renewed interest in the development of BP-based imaging agents due to the increased sensitivity and spatial resolution of this imaging technique. In recent years, a flurry of new BP tracers, many using the generator-produced positron-emitting isotope gallium-68 (t_1/2_ = 68 min), have been reported, mainly focusing on applications for the detection of bone metastases [6-23]. However, the current most-used tracer for the PET imaging of bone metastases in the clinic is [^18^F]NaF, which functions due to the fluoride ion’s ability to displace the hydroxyl group in the structure of HAp (Figure 1a) [24-26].

**Figure 1.**
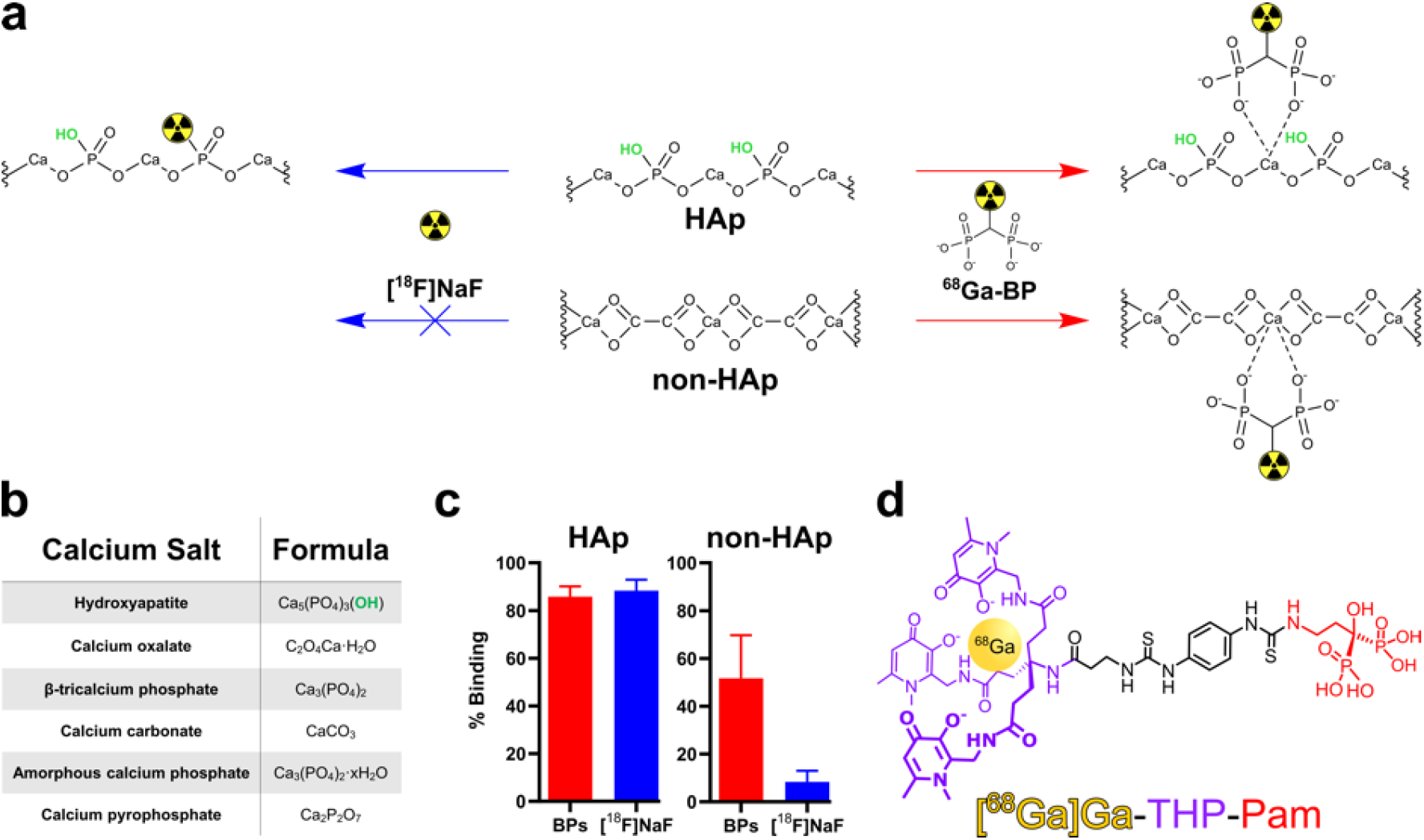
(a) Schematic showing the different binding modes of [^18^F]NaF and a generic BP to calcium salts, including the lack of reaction between [^18^F]NaF and salts without hydroxyl anions. (b) Formulae of selected biologically relevant calcium salts. (c) Summary of *in vitro* binding data of selected ^68^Ga-BPs presented in our previous work [23] in comparison to [^18^F]NaF in HAp and the non-HAp salts listed in panel B. (d) Structure of [^68^Ga]Ga-THP-Pam with the radiometal, gallium-68 shown in yellow; the chelator, THP, shown in purple; and the calcium-binding group, pamidronate, shown in red.

Previously, we have demonstrated that the binding mode of [^18^F]NaF leads to binding specificity towards HAp over other calcium minerals (Figure 1a–c) [23]. However, BPs - which bind through the interaction of the phosphonate groups with calcium ions - have a much broader range of calcium mineral affinity [3, 23]. The prevalence of HAp in bones makes [^18^F]NaF an excellent tracer for the imaging of bones. On the other hand, reports on the mineral composition of extraosseous calcification (EC), which is defined as “deposition of calcium in tissues outside of bone” [27] and includes conditions such as vascular calcification, are diverse and contradictory [28-41]. For example, often these calcifications are euphemistically called HAp as a blanket term for all forms of apatite or solid calcium-containing mineral. However, the general consensus in the literature is that early calcification begins as microcalcifications of amorphous calcium phosphate and whitlockite and, as the calcification progresses, HAp crystals develop and merge into larger sheets or plaques of macrocalcification [42]. However, this is dependent on a number of factors including the location of the calcification and its underlying cause [43]. This consensus is oversimplified and a range of calcium salts may be observed including HAp, apatite, (amorphous) calcium phosphate, whitlockite, calcium oxalate and calcium carbonate [43]. Various studies have demonstrated the presence of calcification in a wide variety of diseases, including atherosclerosis [32, 33, 37-39, 44, 45], age-related macular degeneration [46], Alzheimer’s disease [47], muscular dystrophy [30], various cancers [28, 48-56], kidney stones [34, 35, 40, 57] and chronic kidney disease (CKD) [58, 59].

Nonetheless, [^18^F]NaF is the only clinically used PET tracer for the imaging of extraosseous calcification [60]. Given the potentially diverse forms of calcium mineral present in such calcification, we hypothesised that a HAp-selective tracer such as [^18^F]NaF is not the most appropriate imaging agent for a condition in which HAp may not be the most common form of calcium mineral. To test this hypothesis, we have compared the performance of [^18^F]NaF with the BP-based conjugate of the ^68^Ga chelator tris(hydroxypyridinone) (THP)—[^68^Ga]Ga-THP-Pam (Figure 1d)—a PET tracer that we have recently reported [23], using a rat model that develops macro- and microcalcification across several major organs, and we then highlight the differences in imaging results that stem from the mineral composition of the calcification.

## Materials and Methods

### Materials

All chemicals were purchased from commercial sources unless stated otherwise. Gallium-68 was eluted as [^68^Ga]GaCl_3_ from an Eckert & Ziegler (Germany) ^68^Ge/^68^Ga generator in ultrapure HCl (5 mL, 0.1 M) manufactured to good manufacturing practice (GMP) requirements (ABX, Germany). [^18^F]NaF in H_2_O was purchased from Alliance Medical, UK.

### Synthesis of THP-Pam

THP-Pam was synthesised using an adapted version of our previously published method [23]. In brief, THP-NCS (10.0 mg, 10.4 μmol, CheMatech, France) was dissolved in water (400 μL) and triethylamine (42.4 μL, 30.8 mg, 0.304 mmol, Merck Life Science UK Limited, UK). Pamidronate disodium (29.0 mg, 104 μmol, synthesised as previously published [23]) was dissolved in water (400 μL) and triethylamine (42.4 μL, 30.8 mg, 0.304 mmol, Merck Life Science UK Limited, UK). The THP-NCS solution was added to the pamidronate solution and stirred at 90 °C for 3 h. The crude solution was purified by semi-preparative HPLC: column: Agilent ZORBAX Eclipse XDB-C18 (9.4 × 250 mm, 5 μm); solvent A: water + 0.1% formic acid; solvent B: acetonitrile + 0.1% formic acid; flow rate = 4 mL min^−1^; 0–5 min = 5% B, 5–40 min = 5–20% B, 40–41 min = 20–95% B, 41–45 min = 95% B, 45–46 min = 95–5% B, 46–50 min = 5% B. The identity of the product was confirmed by LC/MS, ^1^H NMR and ^31^P NMR.

### Radiochemistry

[^68^Ga]Ga-THP-Pam was synthesised using our previously published method [23], giving a product with a concentration of 82.4–176.0 MBq mL^−1^ at end of synthesis, and filtered through a 0.22 μm syringe filter prior to injection into animals.

[^18^F]NaF was diluted to a concentration of 115.1–136.3 MBq mL^−1^ at the time of dilution using 0.9% saline and filtered through a 0.22 μm syringe filter prior to injection into animals.

### Rat model of extraosseous calcification

Sprague Dawley rats (male, aged 21–27 days on arrival) were acclimatised for 7 days. The rats were then fed a specialised diet (manufactured by LBS Biotechnology, UK, supplied by Special Diet Services, UK) for 12 days (days 0–11 inclusive). The diet consists of a vitamin K-deficient normal rodent diet supplemented with warfarin (3 mg g^−1^ food, TCI UK Ltd, UK) and vitamin K_1_ (1.5 mg g^−1^ food, Apollo Scientific, UK). For the final four days of the diet (days 8–11 inclusive), each rat was injected subcutaneously with cholecalciferol (vitamin D_3_, 5 mg kg^−1^ day^−1^). The stock solution of cholecalciferol was prepared by addition of cholecalciferol (33.0 mg, 85.8 μmol, Sigma-Aldrich, UK) to absolute ethanol (200 μL) and Kolliphor EL (1.4 mL, Sigma-Aldrich, UK), which was mixed in the dark for 15 min. D-(+)-glucose (750 mg, 4.16 mmol, Sigma-Aldrich, UK) was dissolved in water (18.4 mL) and added to the cholecalciferol solution and mixed in the dark for 15 min. The stock solution was stored in the dark at 4 °C for up to 3 days. Rats were returned to a healthy diet at the end of day 11. Control rats were Sprague-Dawley rats (male, aged 21–48 days upon arrival) which were fed a healthy diet and not injected with cholecalciferol.

### PET/CT imaging and biodistribution studies

On day 11, rats were anaesthetised by inhalation of isoflurane (1.5–4% in oxygen) and the tail vein was cannulated using saline. Each rat was injected intravenously with [^68^Ga]Ga-THP-Pam (100 ± 20 μL, 1.0–6.6 MBq). The rat was maintained under anaesthetic on a warm bed to maintain body temperature for 30 min. The rat was placed in a Mediso nanoScan® PET/CT scanner, where anaesthesia was maintained, and the bed was heated to maintain normal body temperature and CT (55 kVp) was performed. At 1 h post-injection, 1 h PET acquisition (3 × 20 min fields of view, 1:5 coincidence mode; 5-ns coincidence time window) was performed. On day 12, the rats were anaesthetised by inhalation of isoflurane (1.5–4% in oxygen) and the tail vein was cannulated. Each rat was injected intravenously with [^18^F]NaF in saline (100 ± 15 μL, 2.7–11.7 MBq) and imaged at 1 h post-injection on the PET/CT scanner using the same procedure used on day 11. At the end of the scan, the animal was culled at 2 h post-injection for *ex vivo* biodistribution studies. Additionally, non-imaging rats were anaesthetised by inhalation of isoflurane (1.5–4% in oxygen) and the tail vein was cannulated. The rats were injected intravenously with [^68^Ga]Ga-THP-Pam (100 ± 5 μL, 3.9– 10.9 MBq). The rats were maintained under anaesthetic on a warm bed to maintain body temperature for 2 h and culled at 2 h post-injection for *ex vivo* biodistribution studies. Organs were harvested, weighed and counted with a gamma counter along with standards prepared from injected material. Organs and vasculature of interest (heart, lungs, stomach, kidneys, aorta, mesenterics and femoral artery) were fixed in 10% neutral-buffered formalin, after weighing, and subsequently embedded in paraffin for further analysis. For autoradiography, sections of abdominal aorta (approximately 1 cm long, centred on the branching points of the celiac and superior mesenteric arteries) from four rats (two fed the EC diet, two fed a healthy diet) were collected 2 h post-injection with the same volume (100 ± 10 μL) of the same stock of [^68^Ga]Ga-THP-Pam during biodistribution studies. The aortas were placed under a PerkinElmer MultiSensitive Phosphor Screen (12.5 × 25.2 cm) for 5 min and the film was transferred to a Typhoon 8600 Variable Mode Imager. The results were processed using the open-source image processing and analysis package Fiji [61]. The image was pseudo-coloured using the mpl-inferno scale.

### μCT-based 3D X-ray histology (XRH)

μCT was performed at the 3D X-ray Histology facility, μ-VIS X-ray Imaging Centre at the University of Southampton (www.xrayhistology.org). 3D X-ray histology (XRH) is a μCT-based imaging technique that allows non-destructive 3D (volume) visualisation of standard formalin-fixed, paraffin-embedded (FFPE) biopsy specimens and can be seamlessly integrated into conventional histology workflows, enabling non-destructive three-dimensional (3D) X-ray histology. FFPE tissue samples were scanned using a custom designed μCT scanner optimised for XRH (based on the Nikon XTH225ST system, Nikon Metrology UK Ltd.). Imaging was conducted at 80 kVp. Some of the imaging parameters such a as the number of projections, voxel size, and the frames per projection were optimised for each organ based on several factors including sample size and position of the tissue on the cassette. Table S4 (supplementary information) lists all critical parameters for each organ. Scanning was conducted on the histology cassette (blocks were not dewaxed), without the addition of X-ray contrast agents to allow for further analysis by means of conventional histology and consequent correlative imaging.

Upon acquisition, the projection data were reconstructed using conventional filtered back projection into 32-bit floating-point volumes using Nikon’s CT reconstruction. Following CT reconstruction, the data volumes were converted to 16-bit and resliced (re-oriented *in silico*) so that a scroll through the stack along the Z-direction emulates the physical histology slicing of the tissue. This way, any XY single slice through the image stack is parallel to the histology cassette and the XY slice-scroll runs from the wax-block’s surface towards the cassette (similarly to the knife on physical sectioning; see supplementary videos) and manually cropped to the boundaries of the tissue within the scan using Fiji [15].

### Histological staining

Histological analysis was performed by IQPath, Department of Neurodegenerative Disease, UCL. Von Kossa staining counterstained with nuclear fast red and Alizarin Red S staining were performed on 5 μm slices of the tissues. Slices were scanned on a Hammatsu S360 digital slide scanner at 40× magnification and visualised using NZConnect software.

### SEM/EDX

For scanning electron microscopy analysis, paraffin wax was removed using pure xylene for two 10-minute intervals. The slides were then mounted on sample holders using double sided carbon adhesive tape, painted with silver conductive paint, and coated with a 5 nm carbon layer. A Hitachi S-3499N at Wolfson Lab in the Archaeology Department of University College London (UCL) and a Zeiss Leo 1525 Gemini at Electron microscope suite in the Materials Science department of Imperial College London were used for imaging. The Energy Dispersive X-ray spectroscopy (EDX) analysis was applied using an Oxford Instrument EDX detector integrated into the microscope. A 10 kV accelerating voltage and 10 mm working distance settings are used to obtain high-resolution images; the secondary electron (SE) mode was used to get topographic information of the samples, while the backscatter electron detector (BSE) mode enables the differentiation of organic and inorganic materials. The Zeiss obtains two SE images at different depths, which helps us to build a better density-dependent colour SEM (DDC-SEM) [62] image of the minerals and the tissue. EDX is performed on selected areas or points to get the elemental composition and distribution.

### Statistics

Statistical analyses were performed using GraphPad Prism 9 (GraphPad Software Inc., USA) software. Data are presented as mean ± 1 standard deviation. Comparisons between results were analysed using unpaired t-tests, with a p-value <0.05 considered statistically significant. Outlying data points in *ex vivo* biodistribution studies may derive from suspected cross contamination of samples during experiments and were excluded using Tukey’s Fences test.

## Results and discussion

[^68^Ga]Ga-THP-Pam is a conjugate, which has not been pharmacologically optimised, of the clinically used BP pamidronate and the gallium chelator tris(hydroxypyridinone) (THP). THP has excellent properties for radiolabelling with gallium-68, which can be achieved quantitatively in 5 min at room temperature at neutral pH [23, 63-68]. Furthermore, THP conjugates can be radiolabelled using a lyophilised kit to which ^68^Ge/^68^Ga generator eluate can be added directly and can be used without the need for further purification, removing many of the typical hurdles on the path to clinical translation [23, 67, 69, 70]. The ability to make [^68^Ga]Ga-THP-Pam using a kit-based synthesis independently of a cyclotron was identified as a clear advantage for clinical use.

[^68^Ga]Ga-THP-Pam and [^18^F]NaF were previously compared, alongside a leading BP-based tracer in research, [^68^Ga]Ga-NO2AP^BP^, in healthy mice [23]. These studies showed that all tracers tested had rapid blood clearance and high bone uptake (with [^18^F]NaF having the highest).

### PET/CT imaging

PET/CT imaging was performed for a period of 1 h, starting 1 h post-injection, with the rats maintained under anaesthesia from before injection until after imaging. This 1 h period prior to imaging was to allow the radiotracers to reach an equilibrium. We demonstrated that the distribution of both drugs had no major changes after 1 h in previous studies with healthy mice [23]. Furthermore, [^18^F]NaF clinical imaging is typically performed 1 h post-injection [71].

Maximum intensity projection (MIP) PET/CT images are shown in Figure 2a. Images with [^68^Ga]Ga-THP-Pam of the rats fed the EC diet (Figure 2a–b) showed bone and kidney uptake and renal/urinary excretion as seen in previous scans of healthy mice [23], but additionally showed significant uptake in the stomach. A similar pattern of uptake has been reported in a previous study of using BP-based tracers to image calcification [72]. The CT images showed an area of high-density tissue colocalised with the PET signal in the stomach (Figure 2b), which was hypothesised to be highly calcified tissue. Similarly, [^18^F]NaF images of rats fed the EC diet showed the expected bone and bladder uptake, but less [^18^F]NaF was retained in the kidneys compared to [^68^Ga]Ga-THP-Pam.

**Figure 2.**
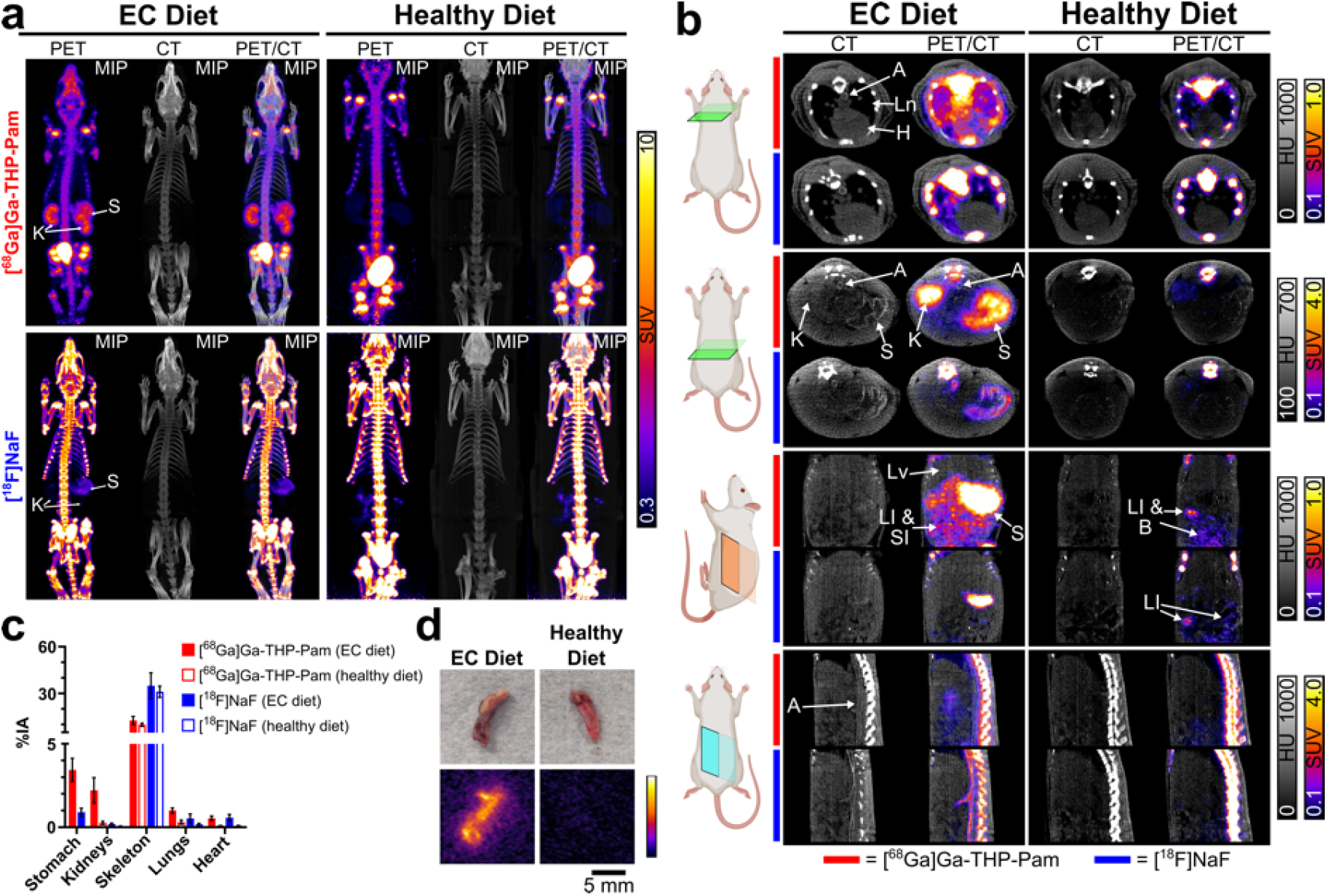
Imaging data of rats with extraosseous calcification and control healthy rats. (a) MIP PET, CT and PET/CT images of rats with extraosseous calcification and control rats with both [^68^Ga]Ga-THP-Pam and [^18^F]NaF 60–120 min post-injection. The images of calcified rats are both of the same animal, and the control images are also data from the same animal. Each PET image shows a standard uptake value (SUV) scale of 0.3– 10. (b) CT and PET/CT images with [^68^Ga]Ga-THP-Pam (in rows indicated by red bars) and [^18^F]NaF (in rows indicated by blue bars) 60–120 min post-injection. The panels show (from top to bottom): axial view of the heart and lungs; axial view of the stomach and kidneys; coronal view of the abdomen; sagittal view of the abdominal spine and aorta. CT and PET scales are matched for each view. (c) quantified PET data of the % injected activity (%IA) in each organ of interest ([^68^Ga]Ga-THP-Pam: n = 4; [^18^F]NaF: n = 3). The entire organ has been included in the ROI, and stomach data from rats fed a healthy diet has been excluded from the data as accurate ROIs could not be drawn due to the stomach not being visible in the images. (d) *Ex vivo* light images (top) and pseudo-coloured autoradiography (bottom) of abdominal sections from rats injected equal volumes of the same batch of [^68^Ga]Ga-THP-Pam and culled 2 h post-injection, showing higher uptake in the aorta of rats fed the EC diet than in rats fed the healthy diet. A = aorta; B = bladder; H = heart; K = kidneys; LI = large intestine; Ln = lungs; Lv = liver; S = stomach; SI = small intestine.

Additionally, [^18^F]NaF showed uptake in the stomach in the rats fed the EC diet, but there was significantly less signal present than in the case of [^68^Ga]Ga-THP-Pam. On the other hand, [^18^F]NaF images of the EC group showed clear uptake in the aorta (Figure 2b), particularly in the abdominal aorta, and major branching arteries, which was not visible in [^68^Ga]Ga-THP-Pam images. Examination of individual slices of the CT images (in the second and fourth rows of Figure 2b) showed clear calcification in the aortas of animals fed the EC diet. The lack of visible [^68^Ga]Ga-THP-Pam uptake is likely due primarily to limitations in the spatial resolution of gallium-68 imaging, particularly evident at the small animal scale, stemming from the high positron energy of gallium-68 (*E*_*avg*_ = 836.0 keV) compared to fluorine-18 (*E*_*avg*_ = 249.3 keV) [73].

The images of the control rats fed a healthy diet demonstrated key differences when compared to rats fed the EC diet. Most evident is the lack of signal in the stomach, both by PET and CT, with both tracers. Also evident is the difference in kidney uptake found in the PET images with [^68^Ga]Ga-THP-Pam, with high signal observed in the EC group and significantly lower signal in the control group. CT imaging, an imaging technique commonly used to image calcification, did not show any areas of hyperintensity in the kidney area that would indicate calcification. Heart and lung uptake of [^68^Ga]Ga-THP-Pam and [^18^F]NaF were also increased in the EC group compared to the control group, but calcification could not be detected by CT. Less prominent increases in uptake of both tracers were observed across several major organs. Single-slice CT and PET/CT images highlighting these differences are shown in Figure 2b.

Quantification of the PET/CT imaging data was performed using regions of interest (ROIs) using the CT as a reference for the skeleton, kidneys, lungs and heart, encompassing the entire organ in each case. Additionally, the PET images were used to generate ROIs for the area of uptake in the stomach in the EC group. This was omitted from the analysis of healthy rats as there was no discernible area visible by PET or CT on which to base an ROI. These data (Figure 2c) indicated an increased accumulation of both [^68^Ga]Ga-THP-Pam and [^18^F]NaF in the kidneys, lungs and heart of the EC group in comparison to the control group. These values are tabulated and p-values are presented in the Supplementary Data (Table S1). Quantification of the PET signal in the stomach of the rats fed the EC diet demonstrated the higher uptake of [^68^Ga]Ga-THP-Pam (3.44 ± 0.69 %IA) than [^18^F]NaF (0.91 ± 0.24 %IA). Comparing the EC and control groups, the largest and most stark difference between the groups was in the kidneys. The kidneys of the EC group showed a mean uptake of 2.21 ± 0.76 and 0.19 ± 0.06 %IA with [^68^Ga]Ga-THP-Pam and [^18^F]NaF, respectively. By comparison, the control group has lower uptake of 0.25 ± 0.13 and 0.07 ± 0.01 %IA with [^68^Ga]Ga-THP-Pam and [^18^F]NaF, respectively. [^68^Ga]Ga-THP-Pam, therefore, showed a mean 8.7-fold (2.02 %IA) increase in kidney uptake in the EC group (compared to the control group) while [^18^F]NaF showed a mean 2.7-fold (0.18 %IA) increase in uptake. Significant increases in uptake were also seen between the EC group and the control group with both tracers in the heart, and with [^68^Ga]Ga-THP-Pam in the lungs. No significant differences were seen in the skeleton with either tracer.

### *Ex vivo* biodistribution

*Ex vivo* biodistribution data at 2 h post-injection (Figure 3) indicated similar trends. Due to the weight loss seen in rats fed the EC diet, and some of the control rats being up to 4 weeks older, data have been presented as standardised uptake value (SUV). The unadjusted data are available in the Supplementary Materials (Table S2).

**Figure 3.**
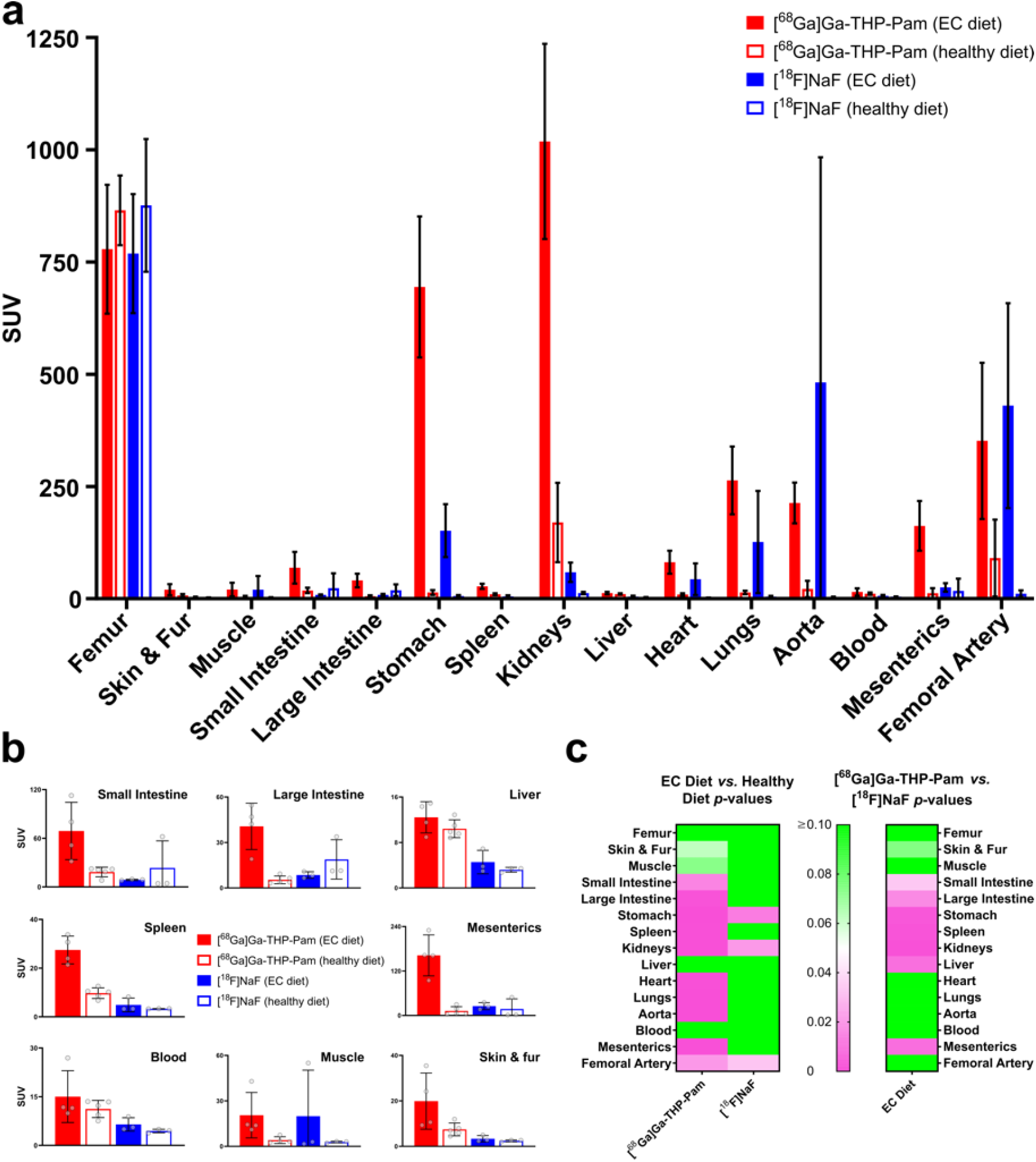
(a) *Ex vivo* biodistribution 2 h post-injection of [^68^Ga]Ga-THP-Pam (EC diet: n = 4; healthy diet: n = 5) and [^18^F]NaF (both diets: n = 3) in rats fed a diet to induce EC and rats fed a healthy diet. (b) Re-scaled *ex vivo* biodistribution in individual organs from panel A, presented for clarity. (c) Heat map of *p*-values resulting from unpaired t-tests when comparing the *ex vivo* biodistribution data of the EC group with the control group with each tracer (left panel) and comparing the two tracers in the EC diet (right panel). The scale has been set to show *p* = 0.05 as white, *p* > 0.05 as green and *p* < 0.05 as pink.

These data largely agree with the trends seen in the images. Both tracers had high uptake in the kidneys and stomach of the EC groups compared to the healthy groups, with [^68^Ga]Ga-THP-Pam showing greater uptake than [^18^F]NaF. Similar trends were observed to a lesser extent in several other major organs, with [^68^Ga]Ga-THP-Pam uptake in organs of rats fed the EC diet being significantly higher than in those of rats fed the healthy diet. The heat map of p-values from unpaired t-tests (Figure 3c) shows the significant increase in [^68^Ga]Ga-THP-Pam uptake in the majority of internal organs. Notably, the aorta signal is greatly increased in the EC group with both tracers, although highly variable in the case of [^18^F]NaF, which has led to a non-significant increase in uptake. This supports the hypothesis that the cause of the aorta not being visible in the [^68^Ga]Ga-THP-Pam images was the comparatively poor spatial resolution of gallium-68 PET images and the proximity to other areas of high activity, including the spine and kidneys. This was further confirmed using autoradiography, in which sections of the abdominal aorta (a few millimetres either side of the branching points of the celiac and superior mesenteric arteries) from rats injected with the same quantity and activity of [^68^Ga]Ga-THP-Pam were extracted and imaged side-by-side on a Phosphor screen. The aortic sections from the animals fed the EC diet were distinctly brighter than the sections from healthy animals (Figure 2d). With this confirmation that [^68^Ga]Ga-THP-Pam is taken up by the aorta of the EC group rats, it should be noted that human anatomy is considerably larger and so partial volume effects are likely to be less of a limiting factor in clinical imaging than in preclinical imaging.

It should also be noted that the high uptake of [^18^F]NaF in the small intestines and large intestines of the healthy group was quite variable. The single instances of high uptake in each organ both occurred in the same rat and the high uptake was also seen in the PET images, therefore these values were not considered to be in any way erroneous. The intestinal uptake of [^18^F]NaF shows high intersubject variability, which we observed in previous healthy animal studies with [^18^F]NaF and is also observed in the clinic, although evidence is anecdotal and the mechanism of uptake remains unknown [74].

### XRH analysis

The data presented so far have shown increased uptake of [^68^Ga]Ga-THP-Pam and [^18^F]NaF across several major organs in the animals fed the EC diet. While calcification in the rats fed the EC diet was evident from the CT in both the aorta and stomach, it was not visible in other organs and hence the cause of the increased uptake of each tracer could not be determined to be calcification without further study. Thus, organs of interest were preserved for histological analysis to determine the extent to which calcification was present. Conventional histological methods for detecting calcification included taking thin (5 μm) slices of tissue and staining using Alizarin Red or the von Kossa stain. This gives a snapshot of calcification that is assumed to be representative of the whole organ. However, this can over-or underestimate the extent of the calcification present in the tissue, or in extreme cases, miss it entirely. For this reason, prior to conventional histology, we scanned the FFPE organs using μCT-based 3D X-ray histology (XRH). This technique allowed for detailed visualisation of the extent of calcification in a whole organ [75]. The high spatial resolution (15–25 μm, dependent on sample) allowed for the detection of calcifications smaller than those able to be detected by preclinical CT, providing an informative intermediate step between preclinical imaging and conventional histology. Animated scrolling MIPs and photorealistic renderings of the XRH data are available online (Supplementary information Table S3). XRH was followed by conventional histology with Alizarin Red (which stains calcium red) and von Kossa (which stains phosphate brown) staining to further confirm that areas identified by XRH as calcification were so. The results of these combined histological techniques (Figure 4) confirmed the presence of calcium deposits in the stomach, aorta, kidneys, small intestine/mesenterics, heart, lungs and aorta. Of note is the localised nature of the calcification in the heart, which is absent from the myocardium and situated solely within part of the aorta collected when excising the heart, which meant that calcification was not evident from the conventional histology of the myocardium slices used for staining.

**Figure 4.**
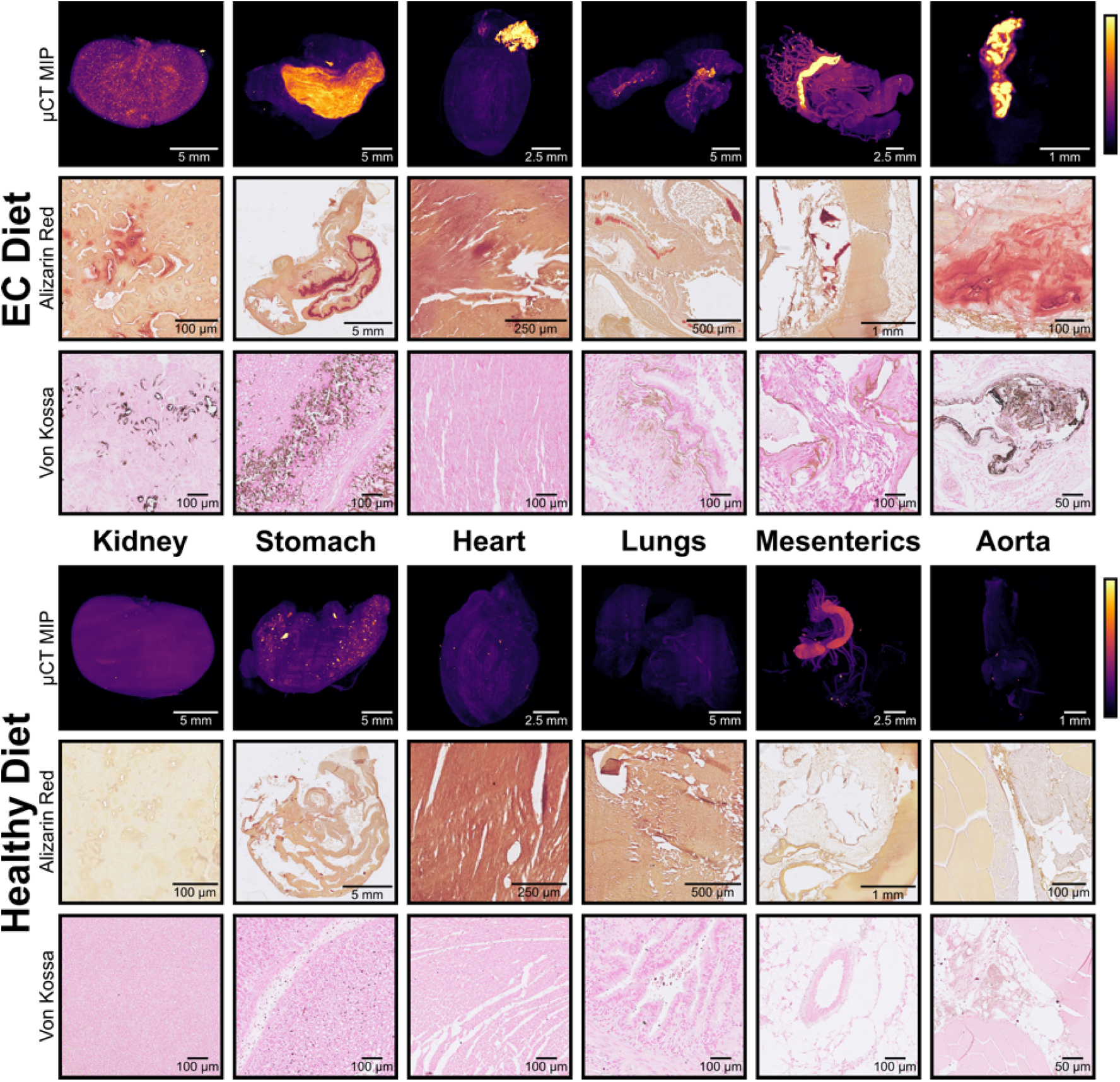
*Ex vivo* analysis of organs to identify calcification. The top half shows organs from rats fed the EC diet, the bottom half shows organs from rats fed the healthy diet. Each row is labelled with its modality. μCT MIP images show pseudo-coloured whole-organ images, both images for each organ have been calibrated to the same colour scale using the air and paraffin wax as reference points. Alizarin Red and von Kossa images show representative 5 μm slices from the organ stained with Alizarin Red S and von Kossa stains respectively to detect calcium. Both images for each organ are shown at the same scale.

### Analysis of mineral composition

Following the identification of calcification in several organs by XRH and histology, the elemental composition and morphology of these deposits were investigated at higher spatial resolutions using SEM and EDX spectroscopy. Comparisons were made between the aorta— where the differences between [^68^Ga]Ga-THP-Pam and [^18^F]NaF uptake were small—and the stomach and kidneys, where differences in uptake were much greater.

SEM images indicated a variety of morphologies which varied by organ. In the kidneys (Figure 5a–c), solid calcium mineral was observed to consist of many small calcifications, in agreement with the μCT data. Each calcification nodule in the kidney has a porous appearance (Figure 5b), which is typical of earlier stage calcification with a less crystalline structure and can likely be considered amorphous calcium phosphate or whitlockite. This calcification was evident from the [^68^Ga]Ga-THP-Pam images but [^18^F]NaF uptake was less clear (Figure 2a). By contrast, the SEM images of the aorta (Figure 5d–f) show a singular solid plaque with a slab-like appearance. This is much more typical of more advanced disease and cardiac calcification, likely to be HAp, which forms as the plaque stabilises and becomes less prone to rupture [42, 76]. Both tracers showed increased uptake to a similar degree in this calcification in the aorta, but [^18^F]NaF was more evident from images due to the different spatial resolutions of the radionuclides as previously discussed.

**Figure 5.**
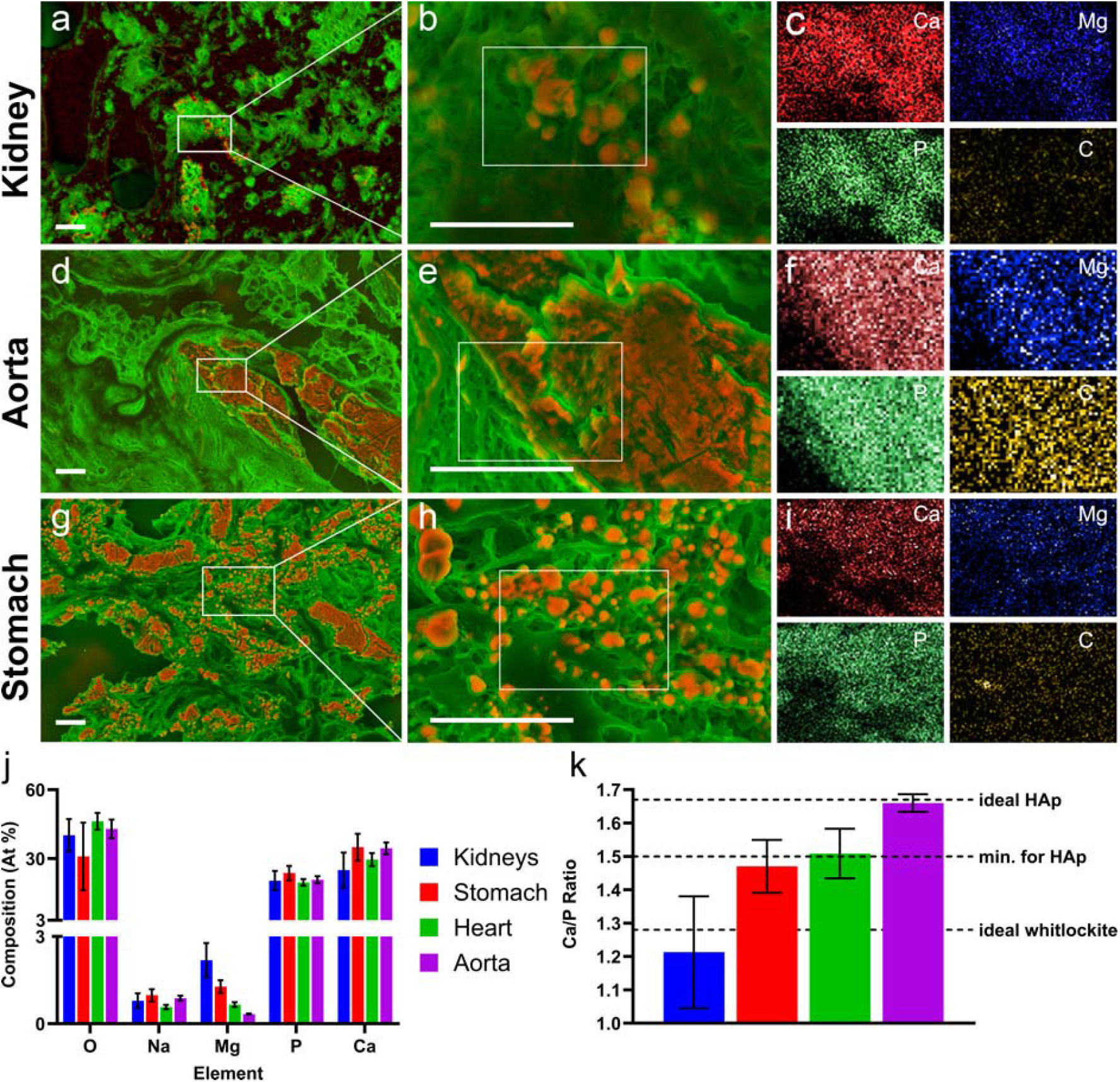
Density-dependent colour SEM (DDC-SEM) [62] EDX images (panels a, d, g: red/orange dense material (mineral) and green less dense material (organic compounds)) and zoom (panels b, e, h) with calcium-rich mineral shown in red/orange and elemental maps (multi-panels c, f, i) for calcium (red), magnesium (blue), phosphorus (green) and carbon (yellow). (a–c) Kidneys. (d–f) Aorta. (g–i) stomach. (j) EDX results of atomic composition of calcifications found in the kidneys, heart, stomach and aorta. (k) Molar calcium-to-phosphorus ratios of mineral present in each organ as determined by SEM/EDX. Data are not absolutely quantitative and should be interpreted only as an indication. The ideal stoichiometric Ca/P ratio of HAp, the minimum ratio required to begin the formation of HAp [77] and the ideal stoichiometric Ca/P ratio [78] are indicated in panel k.

Finally, the stomach (Figure 5g–i) shows a mixture of these sorts of morphologies. Figure 5g shows larger slab-like plaques, while the zoomed-in section (Figure 5h) shows smaller nodules. Figure 5h shows two subtly different morphologies. The centre of the image shows smaller irregularly shaped calcifications similar to those in Figure 5b. These are typical of whitlockite and unlikely to change as disease advances. On the left of the image, larger, more spherical nodules can be seen. These are typical of early-stage HAp development, and these are likely to grow and merge (two can be seen merging in the top left of the panel) into larger slab-like plaques as the disease progresses. Both [^68^Ga]Ga-THP-Pam and [^18^F]NaF demonstrated clearly increased uptake in calcified stomachs, but with [^68^Ga]Ga-THP-Pam having a larger effect.

The elemental mapping in Figure 5c, f and i is shown relative to the white box shown in the zoomed-in panels (b, e and h). These show high levels of calcium corresponding to visible areas of calcification in panels b, e and h. Furthermore, increased levels of phosphorus relative to carbon are also shown in these figures. This indicates the presence of calcium phosphate, which could be present as HAp or another form of calcium phosphate. Figure 5c also shows increased levels of magnesium relative to carbon. The presence of Mg^2+^ cations has been reported to inhibit the formation of HAp, stabilising amorphous calcium phosphate and whitlockite [31, 34, 35, 78, 79], which has been identified in the arteries of CKD patients [40].

These data were further analysed for the atomic composition of the calcifications found within each of the organs, these results are shown in Figure 5j–k. Elemental analysis with EDX is semi-quantitative so these data are interpreted only as approximations and not as definitive. Nonetheless, interesting differences between the organs were observed. Firstly, the Ca/P ratios of the stomach (1.47 ± 0.06) and kidneys (1.21 ± 0.16) were not consistent with the theoretical Ca/P ratio of HAp (1.67) and, in the case of the kidneys, the Ca/P ratio was significantly below the reported minimum Ca/P ratio at which HAp is present in biological mineral (1.50) [77]. The aorta however, showed a Ca/P ratio (1.66 ± 0.02) matching the theoretical Ca/P of Hap and in agreement with the highly crystalline structure observed in Figure 5d–e. Secondly, the levels of magnesium present in the mineral were significantly higher in both the kidneys and stomach than in the heart and aorta. Magnesium is present in whitlockite (Ca_18_Mg_2_(HPO_4_)_2_(PO_4_)_12_, theoretical Ca/P ratio 1.28), which has been identified as the second most abundant mineral in bone [78], although this has been disputed [80]. Indeed, the dispute claimed that whitlockite is present almost exclusively in pathological calcification and teeth, with high levels detected in calcification of several major arteries [80]. Further in-depth analysis of calcification composition was not performed, however, there is a gap in the literature for wide-ranging studies into the composition of calcifications from different stages of calcification and with differing underlying pathologies which have been excised from human patients.

The observed uptake with each tracer in the stomach in combination with the SEM and EDX data and our previous *in vitro* data are consistent with the hypothesis that the composition of calcification is diverse, and that [^18^F]NaF—which almost exclusively targets HAp—may not be as well-suited to the imaging of extraosseous calcification as it is to bone. Furthermore, taking into account the reported trend for HAp to form at later stages of calcification, while amorphous calcium phosphate and whitlockite are more prevalent at earlier stages, BP-based imaging agents may not only offer increased sensitivity at later timepoints but potentially even greater increased sensitivity at earlier timepoints of the EC process compared to both CT and [^18^F]NaF PET. Further studies to investigate this hypothesis should focus on the stage of calcification as well as the characterisation of calcification at different stages in various diseases.

## Conclusions

In summary, we have compared the PET imaging performance of a ^68^Ga-labelled BP PET radiotracer—[^68^Ga]Ga-THP-Pam—to the leading clinical PET imaging option—[^18^F]NaF—in a model of EC in rats. The results showed a significantly increased uptake of both tracers in several major organs in the EC group rats, especially in the stomach and kidneys, with the increase in uptake being greater with [^68^Ga]Ga-THP-Pam than with [^18^F]NaF. The presence of calcification in these organs was confirmed by XRH and conventional histological staining. Finally, the composition and morphology of the calcification were studied by SEM and EDX, which demonstrated that in organs where both [^68^Ga]Ga-THP-Pam and [^18^F]NaF performed similarly by PET, the composition more closely matched theoretical HAp. Yet, in organs in which [^68^Ga]Ga-THP-Pam detected calcification more sensitively than [^18^F]NaF, the composition varied away from theoretical HAp. Therefore, we propose that BP-based tracers may be more appropriate for imaging EC than the HAp-specific [^18^F]NaF, offering the possibility of earlier detection than [^18^F]NaF and CT, before mineral matures into HAp and reaches sufficient density for CT detection.

## Acknowledgements

This work was funded by the EPSRC Centre for Doctoral Training in Medical Imaging [EP/L015226/1], Theragnostics Ltd., the Wellcome/EPSRC Centre for Medical Engineering [WT/203148/Z/16/Z], and the EPSRC programme for next generation molecular imaging and therapy with radionuclides [EP/S032789/1]. Further support comes from a Wellcome Trust Multi User Equipment Grant: A multiuser radioanalytical facility for molecular imaging and radionuclide therapy research [212885/Z/18/Z] and the National Institute for Health Research (NIHR) Biomedical Research Centre based at Guy’s and St Thomas’ NHS Foundation Trust and KCL [grant number IS-BRC-1215-20006]. PET and SPECT scanning equipment at KCL was funded by an equipment grant from the Wellcome Trust under grant number [WT 084052/Z/07/Z]. The authors acknowledge the μ-VIS X-ray Imaging Centre (supported by EPSRC grant EP/H01506X/1) and the Biomedical Imaging Unit at the University of Southampton, for the provision of imaging, data processing and management infrastructure and expertise. This work was supported by the Wellcome Trust Biomedical Resource and Technology Development Grant 212940/Z/18/Z. The views expressed are those of the authors and not necessarily those of the NHS, the NIHR or the Department of Health. This research was funded in whole, or in part, by the Wellcome Trust [WT 203148/Z/16/Z] [212885/Z/18/Z] [WT 084052/Z/07/Z]. For the purpose of open access, the author has applied a CC BY public copyright licence to any Author Accepted Manuscript version arising from this submission.

