## Supplemental information for "^68^Ga-Bisphosphonates for the Imaging of Extraosseous Calcification by Positron Emission Tomography"

| %IA | [$^{68}$Ga]Ga-THP-Pam | | | | [$^{18}$F]NaF | | |
|---|---|---|---|---|---|---|---|
| | VC Diet (n = 4) | | | Healthy Diet (n = 4) | VC Diet (n = 3) | | Healthy Diet (n = 3) |
| | mean ± SD | p-value A | p-value B | mean ± SD | mean ± SD | p-value C | mean ± SD |
| **Stomach/Tissue** | 3.44 ± 0.69 | N/A | ** | N/A | 0.91 ± 0.24 | N/A | N/A |
| **Left Kidney** | 2.15 ± 0.57 | *** | ** | 0.29 ± 0.20 | 0.19 ± 0.07 | * | 0.07 ± 0.02 |
| **Right Kidney** | 2.28 ± 0.95 | ** | * | 0.22 ± 0.01 | 0.19 ± 0.05 | * | 0.07 ± 0.00 |
| **Skeleton** | 12.57 ± 2.62 | ns | ** | 9.85 ± 1.04 | 34.95 ± 8.28 | ns | 31.06 ± 3.63 |
| **Lungs** | 1.00 ± 0.15 | *** | * | 0.30 ± 0.13 | 0.53 ± 0.27 | ns | 0.17 ± 0.06 |
| **Heart** | 0.54 ± 0.13 | *** | ns | 0.10 ± 0.00 | 0.58 ± 0.19 | * | 0.09 ± 0.03 |

**Table S1**. Percentage of the injected activity to accumulate in each ROI 60–120 min post-injection, based on the ROI analysis of the PET-CT images. Stomach/Tissue represents the unidentified region of calcified tissue in the vicinity of the stomach and data are not included for control animals due to the absence of this region. Significance was calculated using an unpaired t-test. [A] = p-value with respect to [$^{68}$Ga]Ga-THP-Pam Healthy Diet group. [B] = p-value with respect to [$^{18}$F]NaF VC Diet group. [C] = p-value with respect to [18F]NaF Healthy Diet Group. ns = not significant; * = p ≤ 0.05; ** = p ≤ 0.01; *** = p ≤ 0.001.

| %IA g$^{-1}$ | [$^{68}$Ga]Ga-THP-Pam | | | | [$^{18}$F]NaF | | |
|---|---|---|---|---|---|---|---|
| | VC Diet (n = 4) | | | Healthy Diet (n = 5) | VC Diet (n = 3) | | Healthy Diet (n = 3) |
| | mean ± SD | p-value A | p-value B | mean ± SD | mean ± SD | p-value C | mean ± SD |
| **Femur** | 6.00 ± 0.30 | ** | ns | 3.22 ± 1.09 | 5.48 ± 0.74 | ** | 2.48 ± 0.47 |
| **Skin & Fur** | 0.16 ± 0.11 | * | ns | 0.03 ± 0.01 | 0.02 ± 0.01 | * | 0.01 ± 0.00 |
| **Muscle** | 0.17 ± 0.13 | ns | ns | 0.03 ± 0.03 | 0.14 ± 0.17 | ns | 0.01 ± 0.00 |
| **Small Intestine** | 0.55 ± 0.23 | ** | * | 0.07 ± 0.03 | 0.06 ± 0.00 | ns | 0.06 ± 0.07 |
| **Large Intestine** | 0.34 ± 0.15 | ** | * | 0.02 ± 0.01 | 0.06 ± 0.01 | ns | 0.05 ± 0.03 |
| **Stomach** | 5.36 ± 0.74 | *** | *** | 0.05 ± 0.02 | 1.08 ± 0.33 | * | 0.02 ± 0.01 |
| **Spleen** | 0.23 ± 0.08 | ** | * | 0.03 ± 0.01 | 0.03 ± 0.02 | ns | 0.01 ± 0.00 |
| **Kidneys** | 7.90 ± 1.26 | *** | *** | 0.54 ± 0.19 | 0.42 ± 0.13 | * | 0.03 ± 0.01 |
| **Liver** | 0.10 ± 0.03 | * | * | 0.04 ± 0.01 | 0.03 ± 0.01 | * | 0.01 ± 0.00 |
| **Heart** | 0.68 ± 0.30 | ** | ns | 0.03 ± 0.01 | 0.31 ± 0.20 | ns | 0.01 ± 0.00 |
| **Lungs** | 2.05 ± 0.56 | *** | ns | 0.05 ± 0.01 | 0.89 ± 0.65 | ns | 0.01 ± 0.00 |
| **Aorta** | 1.69 ± 0.45 | *** | ns | 0.08 ± 0.04 | 3.48 ± 3.01 | ns | 0.01 ± 0.00 |
| **Blood** | 0.13 ± 0.08 | ns | ns | 0.04 ± 0.01 | 0.05 ± 0.01 | * | 0.01 ± 0.00 |
| **Mesenterics** | 1.35 ± 0.60 | ** | * | 0.04 ± 0.03 | 0.18 ± 0.05 | ns | 0.05 ± 0.06 |
| **Femoral Artery** | 2.85 ± 1.59 | * | ns | 0.30 ± 0.19 | 3.08 ± 1.36 | * | 0.03 ± 0.02 |
| **Urine** | 72.38 ± 57.02 | ns | ns | 99.60 ± 59.76 | 4.15 ± 3.81 | ns | 12.38 ± 3.78 |
| **Tail** | 1.04 ± 0.22 | ns | ns | 1.01 ± 0.52 | 1.45 ± 0.32 | * | 0.47 ± 0.05 |

**Table S2**. Biodistribution of [$^{68}$Ga]Ga-THP-Pam and [$^{18}$F]NaF in rats fed a diet to induce VC and rats fed a healthy diet. Significance was calculated using an unpaired t-test. $^{A}$ = p-value with respect to [$^{68}$Ga]Ga-THP-Pam Healthy Diet group. $^{B}$ = p-value with respect to [$^{18}$F]NaF VC Diet group. $^{C}$ = p-value with respect to [$^{18}$F]NaF Healthy Diet Group. ns = not significant; * = $p \leq 0.05$; ** = $p \leq 0.01$; *** = $p \leq 0.001$.

| Group | Organ | Organ ID | Video link |
|---|---|---|---|
| EC Diet | Stomach | 4I | https://drive.google.com/file/d/1IxmT_cBESyCWaYWPzEVRMWb7xBx8EBFf/view |
| Healthy Diet | Stomach | 8I | https://drive.google.com/file/d/1IGfP-J0dYKCV0rNjCUuFtAQhxfr0yok3/view |
| EC Diet | Kidney | 4K | https://drive.google.com/file/d/1PJK_Kz3QsTni1Wyb4g6ADxSF0sE5b2-R/view |
| Healthy Diet | Kidney | 8K | https://drive.google.com/file/d/1QzXjxUeLkogXCdAlqV5SeqsMJOUrE9il/view |
| EC Diet | Aorta | 4P | https://drive.google.com/file/d/1LIBH-k7rPSCrALnwewTfOxFmNu4fn-P2/view |
| EC Diet | Mesenterics | 4S | https://drive.google.com/file/d/17PVyM5l4fb2ZlGTfvbgy-c5lnuSQTnzZ/view |
| Healthy Diet | Aorta | 8P | https://drive.google.com/file/d/1QDGZU7zksO_1hfzCJSkU5CAIodNKIpsY/view |
| Healthy Diet | Mesenterics | 8S | https://drive.google.com/file/d/1dgILuM5wyo3Fhs7aKEiYATzyKkLxqJOQ/view |
| EC Diet | Heart | 13N | https://drive.google.com/file/d/1LQQXchbKCZRvKfybKrGREDTnmHuvFNIc/view |
| EC Diet | Lungs | 13O | https://drive.google.com/file/d/1tO6c_NfkZr7uQXTmQI4P98xC8cWopnS7/view |
| Healthy Diet | Heart | 15N | https://drive.google.com/file/d/11dIQLvrA4UerQ7z_5hHPYe5oltpHSG5f/view |
| Healthy Diet | Lungs | 15O | https://drive.google.com/file/d/1nLYIy82ciOVrQNMjIMrAZeXVWpP3ph7U/view |

**Table S3.** Links to videos scrolling through formalin-fixed, paraffin-embedded organs parallel to the histological cassette.

| | Heart | Stomach | Mesenterics | Lung | Kidney | Aorta |
|---|---|---|---|---|---|---|
| **Energy (kVp)** | 80 | 80 | 80 | 80 | 80 | 80 |
| **Power (W)** | 8.48 | 8.48 | 8.48 | 8.48 | 6 | 6 |
| **Voxel edge (mm)** | 0.0072 | 0.0120 | 0.0120 | 0.0120 | 0.0125 | 0.0060 |
| **Projections** | 5001 | 4001 | 4001 | 4001 | 2601 | 2601 |
| **Frames per projection** | 4 | 4 | 4 | 4 | 8 | 8 |
| **Geometric Magnification** | 20.871 | 12.49 | 12.49 | 12.49 | 11.998 | 24.981 |
| **Estimated scan time (hh:mm)** | 02:40 | 02:07 | 02:07 | 02:07 | 03:15 | 03:15 |

**Table S4**. XRH Imaging parameters for each FFPE organ sample.
